# Transcriptional responses of *Solanum lycopersicum* to three distinct parasites reveal host hubs and networks underlying parasitic successes

**DOI:** 10.64898/2026.01.22.701158

**Authors:** Julia Truch, Maëlle Jaouannet, Martine Da Rocha, Emmanuelle Kulhanek-Fontanille, Cyril Van Ghelder, Corinne Rancurel, Olivier Migliore, Arthur Péré, Stéphanie Jaubert, Christine Coustau, Eric Galiana, Bruno Favery

**Affiliations:** Institut Sophia Agrobiotech, UMR 1355, INRAE, Université Côte d’Azur, 400 routes des Chappes, 06903, Sophia Antipolis, France; EMPR PHYBAC 7006, CNRS, INRAE, Université Côte d’Azur, 400 routes des Chappes, 06903, Sophia Antipolis, France

**Keywords:** Comparative transcriptomic meta-analyses, tomato, hub, network, parasitism, plant pathogens, nematode, oomycete, aphid

## Abstract

Crops face attacks from a wide range of pathogens and pests that deploy various parasitic strategies. All aggressors must manipulate major plant functions, such as immune signaling, hormone pathways, metabolism, and development, to successfully establish themselves and complete their life cycle. Here, we compared the tomato plant’s response to three evolutionary distant pathogens (nematodes, aphids and oomycetes) during compatible interactions using a transcriptomic approach. We identified differentially expressed genes and biological processes, and highlighted potential key host hubs associated with successful parasitism. By integrating recent published datasets, we refined our understanding of the global and tissue-specific mechanisms targeted during compatible interactions and, through co-expression analysis, identified clusters showing shared dysregulation patterns enriched in specific GO terms. Finally, model-to-crop translational analysis using the *Arabidopsis* interactome network, repositioned tomato candidates within larger interaction networks and emphasized the key positions occupied by some of them. Identifying these pivotal tomato targets is crucial to decipher processes underlying parasitism and, consequently, offers new opportunities to develop sustainable multi-pathogen control strategies.

## Introduction

Plants are severely challenged by soil-borne and aerial pathogens that threaten crop production worldwide. Cultivated tomato (*Solanum lycopersicum*), the most important vegetable crop by total production and second by harvested area globally (FAOSTAT2022; https://www.fao.org/faostat/en/#home; 186 Mt/year; 5 M of ha of surface harvested), is grown across a wide range of climates and serves as a host for numerous pathogens, including oomycetes, nematodes, and insects. In resistant plants, early detection of the pathogen through basal immunity (pathogen-associated molecular patterns PAMP-triggered immunity, PTI) or later through the effector-triggered immunity (ETI) modulates MAPK signaling, transcription factors (WRKY, ERF, MYB…), epigenetic modifications, hormone imbalance (salicylic acid, jasmonate, auxins…) leading to productions of reactive oxygen species, pathogenesis-related proteins and secondary metabolites (Campos et al., 2021). Thus, compatible interactions between host plants and parasites rely on the avoidance of plant immunity and the reprogramming of the host’s metabolic activity (Jones & Dangl, 2006). Consequently, pathogens hijack plant resources for their own profit, ultimately leading to reduced plant growth, yield losses, and potentially premature plant death (Kong & Yang, 2023).

As endoparasites, oomycetes and root-knot nematodes (RKN) have mastered long-term interactions within the host roots (Blok et al., 2008; Kamoun et al., 2015). On one hand, zoospores of the oomycete *Phytophthora parasitica* encyst and adhere to the plant surface before penetrating intercellularly. In the roots, *P. parasitica* hyphae invaginate host cell membranes, form haustoria and secrete apoplastic molecules, effectors, that promote pathogen development (Attard et al., 2008). At 15 hours post-infection, *P. parasitica* uses appressoria to penetrate host roots during an initial biotrophic phase (Attard et al., 2010). On the other hand, the mobile infective second-stage juveniles of the RKN *Meloidogyne incognita* penetrate host roots, and migrate intercellularly until reaching the central cylinder, where they become sedentary and induce a feeding structure composed of several giant multinucleate cells. At 21 days post infection, giant cells are fully differentiated and function as mature, metabolically active feeding structures that sustain RKN development (Abad et al., 2008; Favery et al., 2020).

In contrast, aphids such as *Macrosiphum euphorbiae* are ectoparasites establishing durable yet transient interactions by feeding on host plants via sap-sucking (Walling, 2008). Aphid colonies develop on the leaf surface and winged adults ensure the dissemination of new colonies. At 24 hours post-infection in a confined space, aphids have induced their feeding and the host displays a peak response (Jaouannet et al., 2015). Similar to *M. incognita*, *M. euphorbiae* uses a stylet to deliver effectors that manipulate host immunity and metabolism, enabling nutrient acquisition and colony establishment (Favery et al., 2020; F. Wang et al., 2021).

Characterizing both the host targets of effectors and the host functions manipulated is essential to decipher the mechanisms underlying parasitic success, and represents a crucial step to develop new strategies to control pathogens. The diversity of life traits is associated with effector multiplicity which, overall, converges to target directly or indirectly fundamental host components. Interestingly, evolutionary distant pathogens deploy effectors that interact with the same host targets (Mukhtar et al., 2011; Carella et al., 2018). Such shared targets are often interconnected key molecules, hubs, whose perturbation can compromise the robustness of the host regulatory networks. In this context, monitoring plant transcriptional activity is a relevant approach to identify key factors and pathways commonly targeted by pathogens. Some independent works individually analyzed the transcriptomic response of *S. lycopersicum* to the parasites used in our study (i.e. *M. incognita*, *M. euphorbiae* and *P. parasitica*) (Coppola et al., 2013; Naveed & Ali, 2018; Mejias et al., 2021). However none of them investigated and integrated all the responses in a cordonnated experimental procedure and single analysis. Comparative transcriptome meta-analyses focusing on a single model plant or crop species exposed to multiple pathogens, are gaining momentum. These studies reanalyze public RNA-seq datasets to generate multi-pathogens plant response profiles (Biniaz et al., 2022; Y. Wang et al., 2022). Conversely to these studies, which rely on conditions defined by the original experiments, our study used the same tomato variety, matching the tomato reference genome for optimal mapping of RNA-seq data, plants grown under the same conditions and carefully chosen infection time points. To this end, we assessed the transcriptomic response of *S. lycopersicum var. ‘Heinz 1706’* infected or not by the root parasites *M. incognita* (nematode) and *P. parasitica* (oomycete) or the leaf parasite *M. euphorbiae* (aphid), using comparative RNA-seq approaches and pathway analyses. Beyond the identification of genes that are strongly and commonly dysregulated upon infection, we implemented a co-expression analysis to pinpoint candidate hubs associated with parasitic success in roots and leaves and to position homologs of these candidate gene products within their interactome network environment.

## Methods

### Material production

Six four-week-old plants of *S. lycopersicum var. ‘Heinz 1706’* per replicate were grown in a growth chamber under a 16-h photoperiod at 24°C and inoculated with 1000 J2s of *M. incognita* or 500,000 zoospores of *P. parasitica* in the soil, or 50 mixed stages *M. euphorbiae*. Root galls resulting from *M. incognita* infection were collected at 21 days post infection, *P. parasitica*-infected roots were harvested 15 hours post infection, and leaves attacked by *M. euphorbiae* were sampled 24 hours post infection. For each condition, equivalent roots or leaves tissues were collected from non-infected plants. Four independent biological replicates were produced per condition.

### RNA extraction

Samples were ground with mortars in liquid nitrogen and transferred to new tubes. Four volumes of extraction buffer (0.1 M Glycine-NaOH pH 9, 0.1 M NaCl, 0.1% SDS and 1% deoxycholic acid sodium salt) were added. Samples were incubated 5 min on ice, centrifuged for 5 min and supernatants were transferred to fresh tubes. One volume of phenol:chloroform:isoamyl alcohol (PCI; 25:24:1, v/v/v) was added. Samples were vortexed for 15 sec, incubated 2 min at room temperature and centrifuged at 15,000 × g for 15 min at 4°C. The aqueous phase was collected and a second extraction with PCI was performed. The aqueous phase was washed twice with one volume of chloroform before been centrifuged at 12,000 × g for 15 min at 4°C. Sodium acetate was added to a final concentration of 0.3 M, followed by one volume of cold 100% isopropanol. Samples were incubated overnight at -20°C and centrifuged at 12,000 × g for 15 min at 4°C. RNA pellets were washed twice with 1 ml of 70% ethanol, centrifuged at 12,000 × g for 5 min at 4°C, air dried for 10 min, resuspended in 100 µl of RNase free water and 33µl of 8M Lithium Chloride and incubated for one hour at -20°C and centrifuged at 12,000 × g for 5 min at 4°C. Pellets were washed twice with 1ml of cold 70% ethanol, air-dried 10 min and resuspended in 25 µl of RNase free water at 55°C for 5 min. gDNA contamination was removed using Qiagen RNeasy Plus kit following the manufacturer’s instructions.

### Library preparation and sequencing

Library preparation and sequencing were performed by Eurofins Genomics Europe Sequencing GmbH. RNA samples were assessed for integrity and quality and strand-specific, directional polyA selected library preparation was performed using NEBNext Ultra II Directional RNA Library Prep Kit for Illumina with TruSeq Adapter sequences. The samples were sequenced on Illumina NovaSeq 6000 platform using 2x150 Sequence mode.

### Raw data analysis

Raw data were analyzed using an in-house Nextflow based pipeline. Briefly, quality controls of raw data were performed using FastQC (fastqc%3A0.11.9--hdfd78af_1) and MultiQC (multiqc%3A1.10.1--pyhdfd78af_1) before and after data cleaning including trimming and filtering for reads of at least 100 bp using FastP (fastp%3A0.23.2--h79da9fb_0). Reads were then mapped with STAR (star%3A2.7.8a--h9ee0642_1) with the options --genomeSAindexNbases 10 for the indexing and --alignEndsType EndToEnd and --twopassMode Basic for the mapping. For each interaction, reads were mapped on a composite genome combining *S. lycopersicum* v4 genome (solgenomics.net) with either the *M. incognita* V3 genome ((Blanc-Mathieu et al., 2017), the *M. euphorbiae* transcriptome (NCBI ref GAOM01000000) or the *P. parasitica* genome GCA_000247585.2 (https://protists.ensembl.org) in order to remove potential pathogen reads. Then, all reads mapping to pathogen genome/transcriptome were removed to conserve only the reads mapping to the *S. lycopersicum* v4 genome. Read counts were obtained with Feature-Counts (subread:2.0.1--hed695b0_0) with the options -t mRNA and -g ID. Differential analyses were performed using DESeq2 (Love et al., 2014) and EdgeR (Robinson et al., 2010). The gene annotation was based on the *S. lycopersicum* annotation ITAG4.1 (solgenomics.net). Principal component analysis (PCA), VENN diagrams and co-expression analyses in roots and leaves were performed using DiCoExpress (Lambert et al., 2020). Graphical representations were performed with ggplot2 (ggplot2_3.5.0). Filtering, normalization and quality control (including PCA analysis) of the dataset were performed using the Quality_Control option with default parameters (CPM_Cutoff=1 and Normalization_Method = “TMM”). Datasets were analyzed using GLM_Contrasts model. DiffAnalysis.edgeR was run with default parameters (Alpha_DiffAnalysis = 0.05, NbGenes_Profiles = 20, NbGenes_Clustering = 50 and p-values were adjusted for multiple testing using the Benjamini-Hochberg (BH) method to control the false discovery rate (FDR)). Venn diagrams were performed using Venn_IntersectUnion option. Co-expression analyses were performed using Coexpression_coseq option based on Gaussian mixture models using default parameters on gene candidates commonly dysregulated in response to roots or leaves pathogens therefore excluding the candidates displaying significantly different responses between the parasites of interest. Singularity prebuilt images were retrieved from the Galaxy Project Singularity Depot (https://depot.galaxyproject.org/singularity/).

### Subtilase and TCP gene family analyses

Fifty-five protein sequences were retrieved from TAIR (Araport11), matching with the 56 subtilases described in A. Figueiredo et al. (2014), with AT3G44262 removed because it is a pseudogene). Similarly, 103 protein sequences annotated as subtilase were extracted from the *S. lycopersicum* protein dataset (ITAG4.1). The 158 sequences were aligned using MUSCLE (v3.8.31) with default settings for highest accuracy. No outgroup has been added as the objective was to position *S. lycopersicum* sequences according to *A. thaliana* sequences and the known subtilase sub-families. The phylogenetic tree was reconstructed using the maximum likelihood method implemented in the PhyML program (v3.1/3.0 aLRT). The WAG substitution model was selected assuming an estimated proportion of invariant sites (of 0.000) and 4 gamma-distributed rate categories to account for rate heterogeneity across sites. The gamma shape parameter was estimated directly from the data (gamma=1.197). Reliability for internal branch was assessed using the aLRT test (SH-Like). Graphical representation and edition of the phylogenetic tree were performed with iTOL v7. Similarly, 24 and 32 protein sequences annotated as TCP transcription factors were retrieved from the *A. thaliana* (Araport11) and the *S. lycopersicum* (ITAG4.1) protein datasets, respectively. These 56 sequences were aligned with MEGA X using MUSCLE. The evolutionary history was inferred using the Neighbor-Joining method. The bootstrap consensus tree was inferred from 1000 replicates. The evolutionary distances were computed using the JTT matrix-based method.

### Gene ontology (GO) and transcription factor analyses

The GO analyses were performed with GOFuncR (GOfuncR_1.22.2) except the co-expression analysis part for which the GO analyses were performed using DiCoExpress using Enrichment option with default parameters (Alpha_Enrichment=0.01) (Lambert et al., 2020). Visualisation was done with ggplot2 (3.5.0). The list of transcription factors was downloaded from the Plant Transcription Factor Database (PlantTFDB v5.0) (https://planttfdb.gao-lab.org/index.php?sp=Sly) (Tian et al., 2019).

### Ortholog identification, effector interactor and interactome network analyses

Protein sequences from *S. lycopersicum* (ITAG4.1), *Arabidopsis thaliana* (Araport11), *Amborella trichopoda* (AMTR1.0), *Brassica napus* (AST_PRJEB5043_v1), *Capsicum annuum* (ASM51225v2), *Coffea arabica* (GCF_003713225.1_Cara_1.0), *Helianthus annuus* (HanXRQr2.0-SUNRISE), *Medicago truncatula* (MedtrA17_4.0), *Nicotiana benthamiana* (Niben261), *Oryza sativa* (IRGSP-1.0), *Prunus persica* (NCBIv2), *Sorghum bicolor* (NCBIv3), *Selaginella moellendorffii* (v1.0), *Solanum tuberosum* (SolTub_3.0), *Vitis vinifera* (PN40024.v4) and *Zea mays* (Zm-B73-REFERENCE-NAM-5.0) were functionally annotated using InterProScan (5.51-85.0) and the ortholog analysis was performed using OrthoFinder (2.5.4). The identification of effectors interacting with candidate homologs was performed using EffectorK (González-Fuente et al., 2020). *A. thaliana* based interactome was generated with BIOGRID-ORGANISM (v4.4.227) using BIOGRID-ORGANISM-Arabidopsis_thaliana_Columbia-4.4.227.psi25 (Stark et al., 2006) and analyzed with Cytoscape (v3.10.1). The Betweenness Centrality for the subnetworks were analyzed using Cytoscape Analyze Network tool without selecting for Directed Graph.

## Results

### Global transcriptomic response of S. lycopersicum to M. incognita, P. parasitica and M. euphorbiae

We performed comparative RNA-seq analyses on *S. lycopersicum* roots inoculated or not with either *M. incognita* or *P. parasitica* and leaves inoculated or not with *M. euphorbiae,* to assess the transcriptional reprogramming occurring in tomato under biotic stresses generated by root and leaf parasites. To be considered as differentially expressed genes, candidates had to display both, an adjusted p-value < 0.05 in DEseq2 analysis and FDR < 0.05 in EdgeR analysis. Thus, we identified 12,231 differentially expressed genes at 21 days post infection with *M. incognita*, 12,018 differentially expressed genes at 15 hours post infection with *P. parasitica* and 11,001 differentially expressed genes at 24 hours post infection with *M. euphorbiae* (Figure 1a and Figure S1). A total of 4,461 differentially expressed genes were identified in response to the three pathogen infections (Figure 1b). Among them, 961 were down-regulated and 645 were up-regulated across all three conditions, whereas 2,855 showed variable dysregulation patterns depending on the specific interaction studied (Figure 1c). To obtain an overview of the biological processes commonly targeted by the three pathogens, we performed GO enrichment analyses on the lists of differentially expressed genes identified. We identified 53 biological processes GO terms shared across the three interactions (Figure 1d). Among them, 45 GO terms were all either over- or under-represented (Figure S2a), whereas eight showed variability depending on the specific interaction (Figure S2b). All over-represented GO terms were related to metabolic processes, while the negative regulation of metabolic processes were among the under-represented GO terms. We then split the list of differentially expressed genes between up- and down-regulated. The GO enrichment analysis on the list of up-regulated genes showed an over representation of the metabolic process in the tree interactions (Figure S2c and d) whereas not GO term was detected as over represented in all tree interactions when focusing on the down-regulated genes (Figure S2e and f). These results are consistent with the enhanced metabolic activity of the host associated with parasite infection.

**Figure 1.**
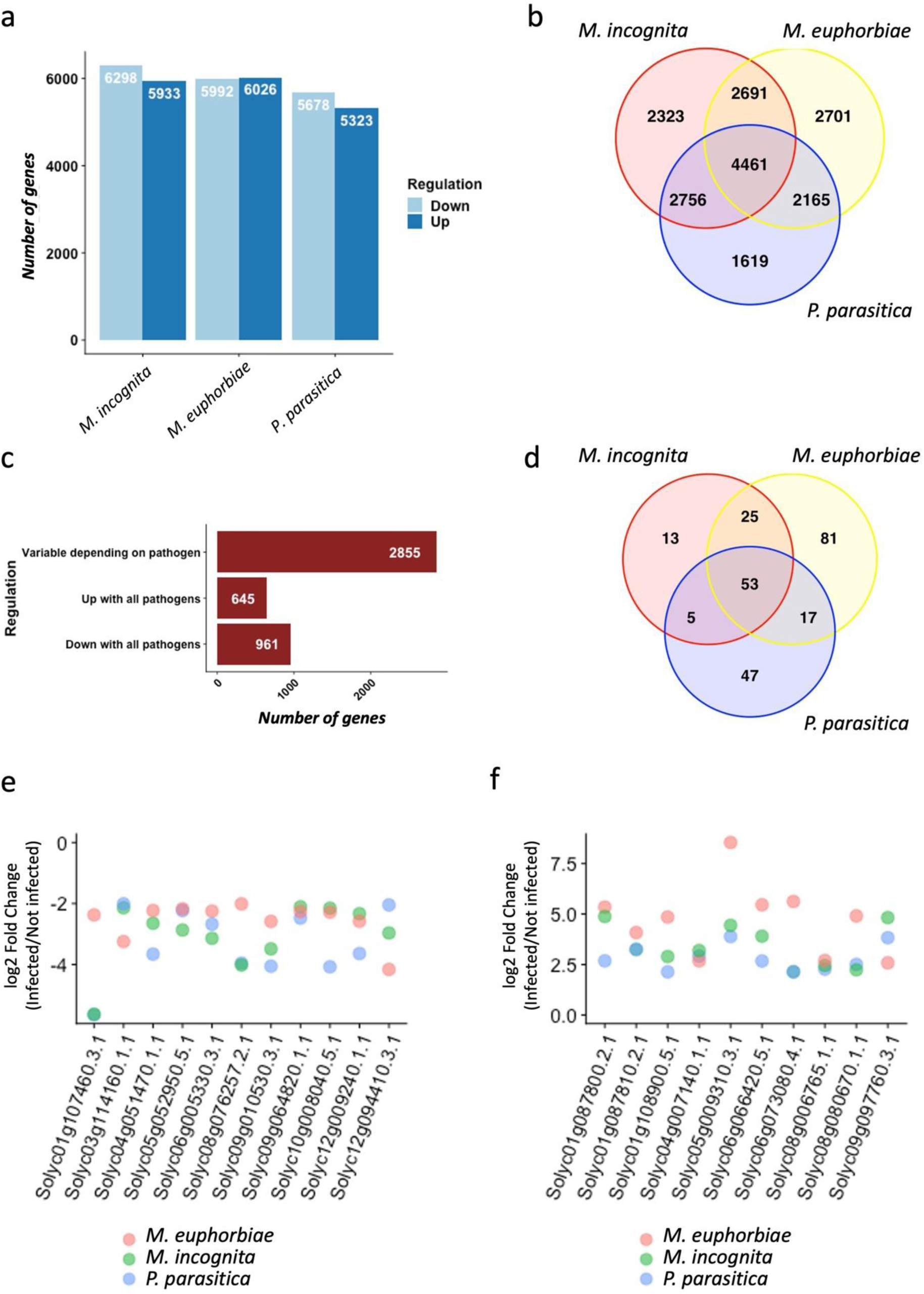
- Transcriptomic response of *S. lycopersicum* to *M. incognita*, *P. parasitica* and *M. euphorbiae*. a. Barplot showing the number of differentially expressed genes up- and down-regulated depending on the tested parasites (log2 Fold Change (Infected/Not infected)). b. Venn diagram illustrating the overlaps between the differentially expressed genes identified in the experiments with the different parasites. c. Barplot comparing the change in gene expression of the 4461 candidate genes differentially expressed during the three interactions. D. Venn Diagrams of GO overlaps between the three interactions considering all the differentially expressed genes. e. Differentially expressed genes with a log2(Infected/Not infected) lower than -2 and an adjusted p-value lower than 0.05 (FDR/Benjamini-Hochberg, DESeq2 (Love et al., 2014)) in the three interactions tested (*M. euphorbiae*, *M. incognita* and *P. parasitica*). f. Differentially expressed genes with a log2(Infected/Not infected) higher than 2 and an adjusted p-value lower than 0.05 (FDR/Benjamini-Hochberg, DESeq2 (Love et al., 2014)) in the three interactions tested, *M. euphorbiae*, *M. incognita* and *P. parasitica* (log2 Fold Change (Infected/Not infected)).

Focusing on the differentially expressed genes most strongly responsive to all three pathogens, we identified only 11 genes with a log2 fold change (log2FC) < -2 across all interactions (Figure 1e). This short list includes the *TOLERANT TO CHILLING AND FREEZING 1* (*AtTCF1*) *Arabidopsis* ortholog, *Solyc05g052950.* This stress response-associated gene encodes for a regulator of chromosome condensation that indirectly affects lignin accumulation (Ji et al., 2015). Surprisingly, the list also contains two *A. thaliana* orthologs of transcription factors that are involved in plant pigment regulation (flavonol and chlorophyll, respectively): the *Solyc06g005330,* related to the AtMYB48 transcription factor and *Solyc12g009240*, a putative ethylene-responsive transcription factor (homolog of ERF017 in *A. thaliana*). Conversely, 10 differentially expressed genes displayed a log2FC > 2 in three interactions (Figure 1f). Among them, we identified genes that are involved in a variety of biological responses. Notably, we identified tomato orthologs of *A. thaliana* proteins known to regulate systemic acquired resistance (SAR), such as the NON-INDUCIBLE IMMUNITY1 (NIM1)-INTERACTING 2-like protein (Solyc04g007140); proteins induced upon biotic stress, such as N-acetyltransferase NATA induced by insect feeding and bacterial infection (Solyc08g006765) (Adio et al., 2011; Lou et al., 2016) or abiotic stresses, such as the chromatin remodeling factor OXIDATIVE STRESS 3 (OXS3, Solyc06g066420) (Blanvillain et al., 2009). Finally, we identified two susceptibility factors: (i) the osmotin-like protein OSM34 (Solyc08g080670), which is subverted in *A. thaliana* by the turnip mosaic virus (TuMV) thus decreases plasmodesmata callose deposition to promote viral cell-to-cell movement and replication (He et al., 2025), and (ii) DOWNY MILDEW RESISTANT6 DMR6, an enzyme that converts SA into its inactive form and whose loss of function in tomato confers broad-spectrum disease resistance (Thomazella et al., 2021). Consistently, both *DMR6-1* (*Solyc03g080190,* Figure S2g) and *DRM6-2* (*Solyc06g073080*, Figure 1f) were upregulated in the three interactions tested.

Furthermore, we identified two subtilases (or subtilisin-like proteases) among the top 10 most strongly upregulated differentially expressed genes in response to the three parasites. Subtilases (SBTs) are serine peptidases involved in general or specific proteolysis, contributing to many molecular processes including growth control, cell wall properties regulation, extracellular signaling molecules, vascular development and programmed cell death (Schaller et al., 2018).These two proteases encoding genes (*Solyc01g087800* and *Solyc01g087810*) were arranged in tandem within a larger subtilase cluster (cluster 1) on the chromosome 1 (Figure 2). The subtilase gene family was particularly well represented in our differentially expressed gene dataset. Indeed, 65 of the 103 annotated subtilase genes in tomato were differentially expressed in response to at least one pathogen. Gene numbers assigned to each SBT family are variable depending on plant species. Our results showed that subtilases in *A*. *thaliana* have more gene members from the families SBT3 and SBT4 than in tomato. Conversely, tomato subtilases undergo a massive expansion in the SBT1 subfamily that includes subtilases from the clusters 1 and 8 (Figure 2). This observation is in line with observations made in *Solanum tuberosum*, another *Solanaceae* species (Schaller et al. 2017). Most of the functional studies related to the SBT1 family are based on P69 protease in tomato (from the cluster 8) which are in the clade of AT1G04110 (SDD1) (Reichardt et al., 2018). P69 paralogs played a role in pathogen defense such as the P69C processing a leucine-rich repeat protein to potentially trigger recognition and initiate immune signaling. P69 are also targeted by different pathogen effectors coding for protease inhibitors. A deeper focus on the subtilase cluster 1 revealed additional subtilase candidates differentially expressed in each plant-parasites interaction (Figure 2). Interestingly, both clusters included subtilases identified as phytaspases, caspases-like proteases regulating programmed cell death, including *Solyc01g087800,* the most highly expressed candidate. The phylogenetic tree showed that subtilases from cluster 1 are all related to the *A. thaliana* gene *SBT1.9* (*At5g67090*), another subclade of the SBT1 group of sequences for which functional studies are required.

**Figure 2.**
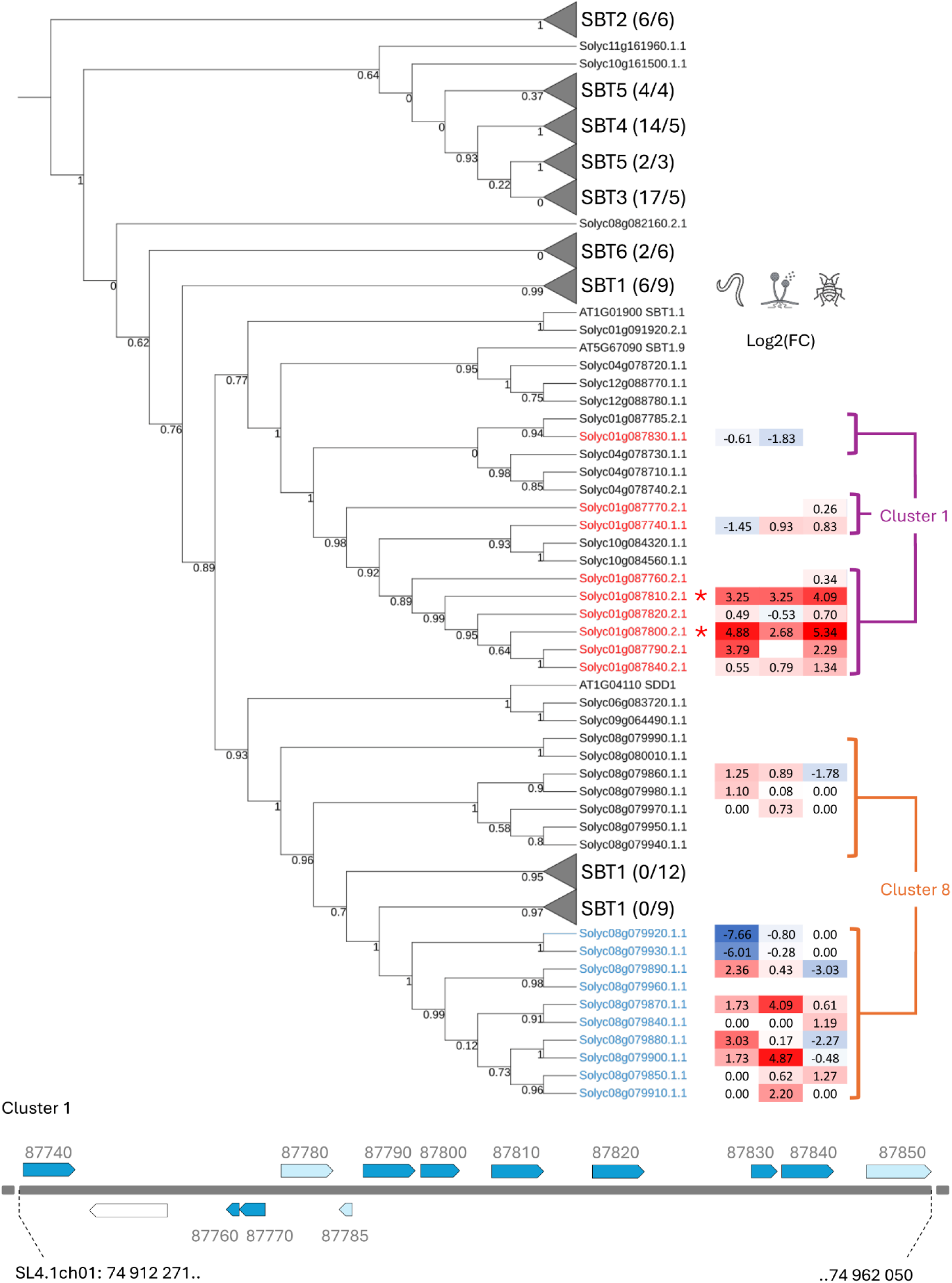
- Phylogenic tree of subtilases of the cluster 1 and 8 in the chromosome 1 and 8 of *S. lycopersicum*. The Maximum Likelihood phylogenetic tree (WAG model, aLRT test) contains 55 subtilases from *A. thaliana* and 103 subtilases from *S. lycopersicum*. SBT1-6 subfamilies were assigned following (A. Figueiredo et al., 2014) and clades were collapsed to facilitate the visualization (the extended developed phylogenetic tree is available in Figure S3). In brackets, the number of *A. thaliana* sequences/number of sequences from *S. lycopersicum*. The differential expression (Log2 Fold Change (Infected/Not infected)) of genes from the cluster 1 and 8 (from chromosome 1 and 8 of *S. lycopersicum*) in response to the three pathogens (*M. incognita*, *P. parasitica* and *M. euphorbiae*) is presented on the right panel (adjusted p-value lower than 0.05 (FDR/Benjamini-Hochberg, DESeq2 (Love et al., 2014)), in blue down regulated genes and in red upregulated genes). Asterisks indicate the two highly overexpressed genes in response to the three pathogens. Physical representation of the Cluster 1, including the coordinates on the *S. lycopersicum* chromosome 1, is presented with differentially expressed subtilase genes in response to at least one parasite in dark blue and light blue for the others.

### In-depth identification of targeted hubs during plant-pathogen interactions

Following the identification of the most strongly dysregulated genes, we next analyzed differentially expressed genes through the lens of common stress-response pathways. We first examined the salicylic acid (SA) pathway and, in addition to *DMR6s* and *NIM1-INTERACTING 2-like,* we identified five differentially expressed genes encoding: two MACPF domain-containing proteins (Solyc01g005220 and Solyc02g077780), the latter being implicated in the negative regulation of the SA-related defense responses (Noutoshi et al., 2006), and three NIM1-like proteins (Solyc07g040690, Solyc02g069310 and Solyc07g044980) (Figure 3a). All of these genes were upregulated in the presence of root pathogens (*M. incognita* and *P. parasitica*), whereas their expression pattern were more variable in response to the leaf parasite *M. euphorbiae*. We, then, identified six candidates associated with auxin response, including two members of the small auxin upregulated RNA (SAUR) family (Solyc01g110907 and Solyc07g042470), known to regulate cell elongation and the activation of plasma membrane H^+^-ATPases (Spartz et al., 2016) (Figure 3b). Finally, we identified 11 differentially expressed genes associated to ethylene response in the three interactions, which held the attention given the pivotal role of ethylene in term of regulation of plant development and stress resilience (Figure 3c). Among these differentially expressed genes, we detected several transcription factor encoding genes, including *Solyc05g052050,* which encodes Pti4, a protein interacting with the resistance Pto kinase protein and cis regulatory elements (Zhou et al., 1997) and *Solyc06g053710,* which encodes the ethylene receptor ETR4 (Figure 3c). Using the Plant Transcription Factor Database (Tian et al., 2019), we further identified 34 up-regulated (Figure 3d) and 92 down-regulated (Figure 3e) genes across the three interactions within the “*Solanum lycopersicum* Transcription Factors database”, covering 29 transcription factor families. The representations between TF families differ between up- and down-regulated candidates (Figure 3d and e). Up-regulated genes in all three interactions showed an enrichment in ERF family (p-value = 0.0207, odds ratio = 4.346768, Fisher’s Exact Test) and WRKY family (p-value = 0. 01532, odds ratio = 7.604422, Fisher’s Exact Test) (Figure 3d) compared to down-regulated candidates (Figure 3e).

**Figure 3.**
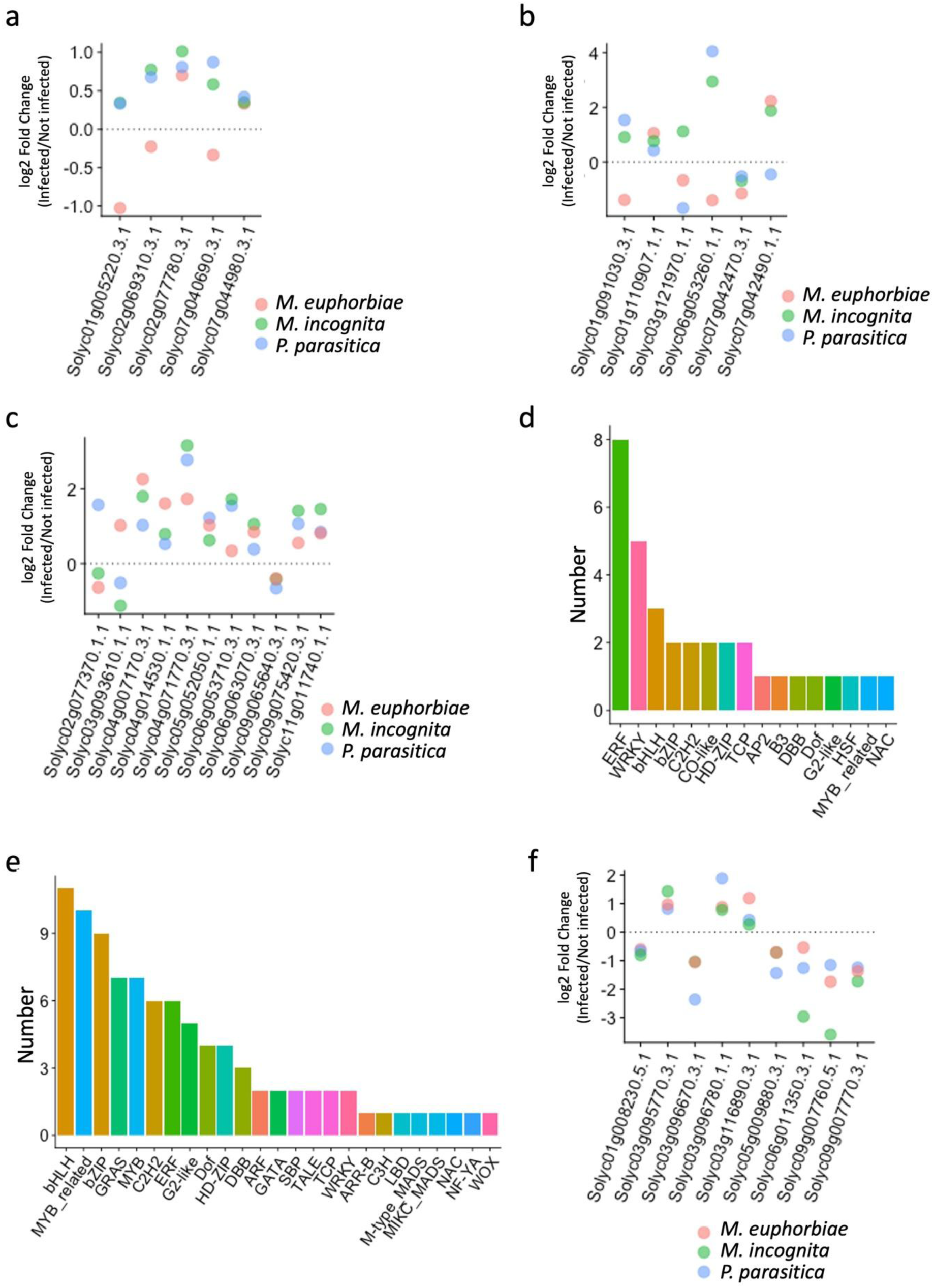
- Characterization of *S. lycopersicum* candidate genes differentially expressed in the three interactions (*M. euphorbiae*, *M. incognita* and *P. parasitica*). a. Differentially expressed genes with an adjusted p-value lower than 0.05 (FDR/Benjamini-Hochberg, DESeq2 (Love et al., 2014)) associated with salicylic acid pathways in the three interactions tested (log2 Fold Change (Infected/Not infected)). b. Differentially expressed genes associated with auxin response in the three interactions tested. c. Differentially expressed genes with an adjusted p-value lower than 0.05 (FDR/Benjamini-Hochberg, DESeq2 (Love et al., 2014)) associated with ethylene response in the three interactions tested (log2 Fold Change (Infected/Not infected)). d. Distribution of candidate genes identified in Plant Transcription Factor Database and up-regulated in the three interactions tested. e. Distribution of candidate genes identified in Plant Transcription Factor Database and down-regulated in the three interactions tested. f. Selected candidate hubs in *S. lycopersicum* encoding for orthologs of effector interactors in *A. thaliana (*adjusted p-value lower than 0.05 - FDR/Benjamini-Hochberg, DESeq2 (Love et al., 2014), (log2 Fold Change (Infected/Not infected)))

We then assessed whether the putative hubs identified by transcriptomics in *S. lycopersicum* might have *A. thaliana* homologs encoding characterized effector hubs, *i.e.* host proteins interacting with several pathogen effectors. We identified orthologs/homologs between *S. lycopersicum* and *A. thaliana* using OrthoFinder (Emms & Kelly, 2019) and search the EffectorK database compiling *A. thaliana* proteins interacting with pathogen effectors (González-Fuente et al., 2020). Among the *S. lycopersicum* orthologs/homologs of *A. thaliana* effector interactors, we identified, among others, candidate hubs encoding WRKYs (*Solyc03g095770* and *Solyc03g116890*), bHLH (*Solyc05g009880*) and TCP (*Solyc01g008230*) transcription factors, a protein phosphatase 2C (PP2CA) (Solyc03g096670), a protein homologous to RESPONSE TO LOW SULFUR 2 (LSU2) (*Solyc03g096780*) and several paralogs of plasma membrane intrinsic proteins (*Solyc06g011350*, *Solyc09g007760* and *Solyc09g007770*) (Figure 3f). Altogether, these results pointed out a short panel of putative key tomato hubs that could be directly or indirectly affected by at least these three pathogens and acting at multiple levels to promote parasitic success.

### Root and leaf parasites trigger preferentially different pathways

To dissect the regulatory networks underlying parasitic success, we performed a co-expression analysis. As our dataset included *S. lycopersicum* root RNA-seq data in response to two telluric parasites (*M. incognita* and *P. parasitica*) and *S. lycopersicum* leaf RNA-seq data in response to only one aerial parasite (*M. euphorbiae*), we expanded our dataset with publicly available *S. lycopersicum* leaf RNA-seq data in response to the South American tomato pinworm *Tuta absoluta* (24 hours post infection) (Roumani et al., 2022) to further explore tissue-dependent co-expression patterns. To ensure consistency across datasets, we reanalyzed the publicly available raw data using the same pipeline and parameters applied to the RNA-seq data from the three other parasites (*M. incognita*, *P. parasitica and M. euphorbiae)*. Principal component analysis (PCA) revealed a clear separation between *S. lycopersicum* leaf and root RNA-seq samples along the first component, accounting for more than 45% of the variance (Figure S4a), consistent with the distinct gene expression patterns observed in roots and leaves (Attard et al., 2010; Pirona et al., 2023). Therefore, we examined the co-expression profiles for root and leaf infections separately using DiCoExpress (Lambert et al., 2020), to identify the genes displaying a common response to the pathogens tested. We thus removed the candidate genes for which the response significantly differs between the parasites tested (*M. Incognita* vs *P. parasitica*, 5,615 genes, Figure 4a and *M. euphorbiae* vs *T. absoluta,* 2,998 genes, Figure 5a).

**Figure 4.**
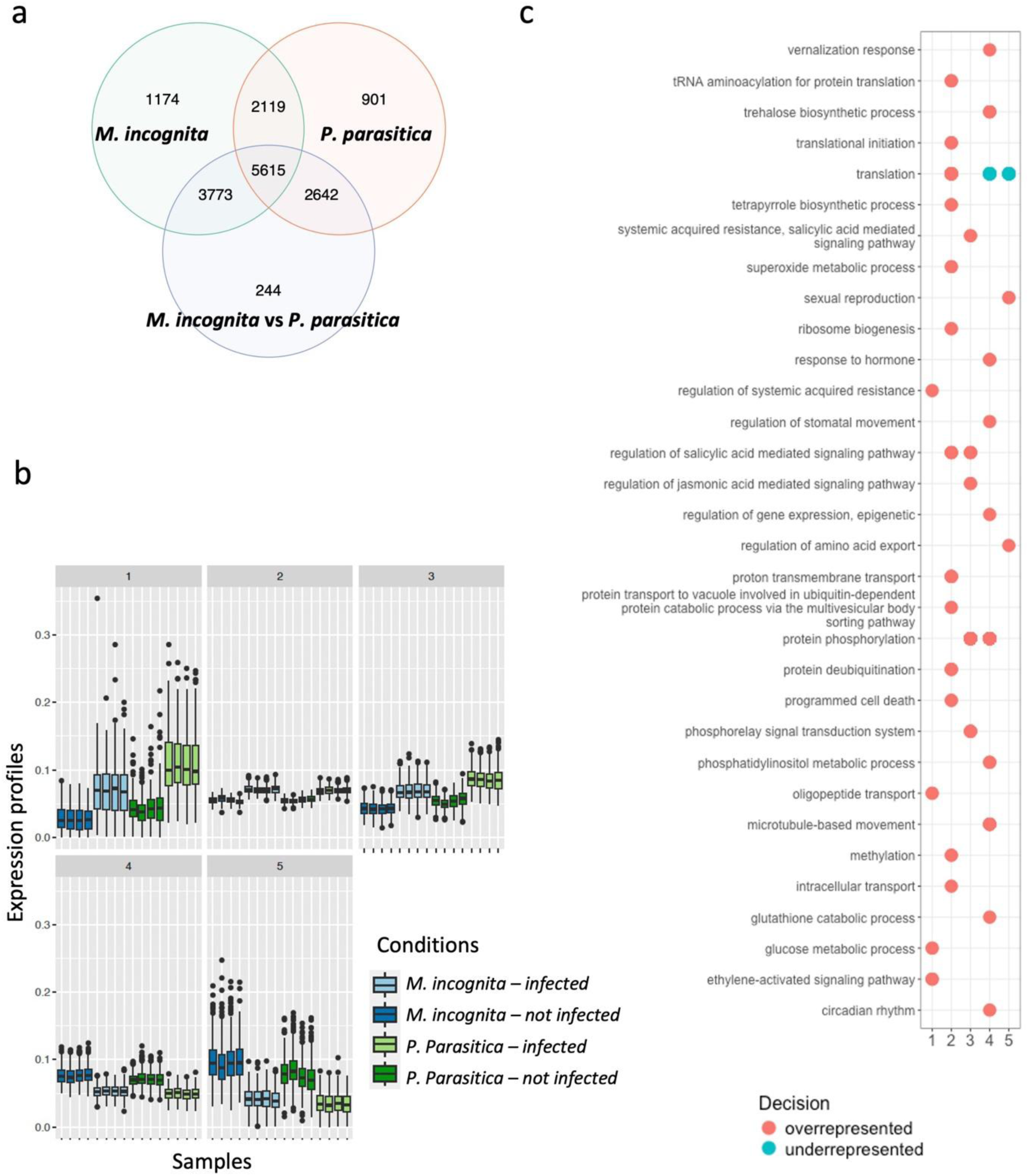
- *S. lycopersicum* root response to telluric parasites (*M. incognita* and *P. parasitica*) revealed 2,119 genes displaying a similar response to both parasites. (a) Venn diagram of differentially expressed genes in roots in response to *M. incognita (in green)* and *P. parasitica (in red) and the genes significantly differing in their response between the two root parasites - M. Incognita vs P. parasitica - (in blue)* based on generalized linear model and likelihood ratio test-based p-values adjusted lower than 0.05 (Benjamini-Hochberg) based on the EdgeR package (Robinson et al., 2010) via DiCoExpress (Lambert et al., 2020). (b) Co-expression showing the five clusters with boxplot expression profiles per sample per conditions. (c) GO enrichment of biological processes for the five co-expression clusters containing gene candidates commonly dysregulated in presence of root pathogens.

**Figure 5.**
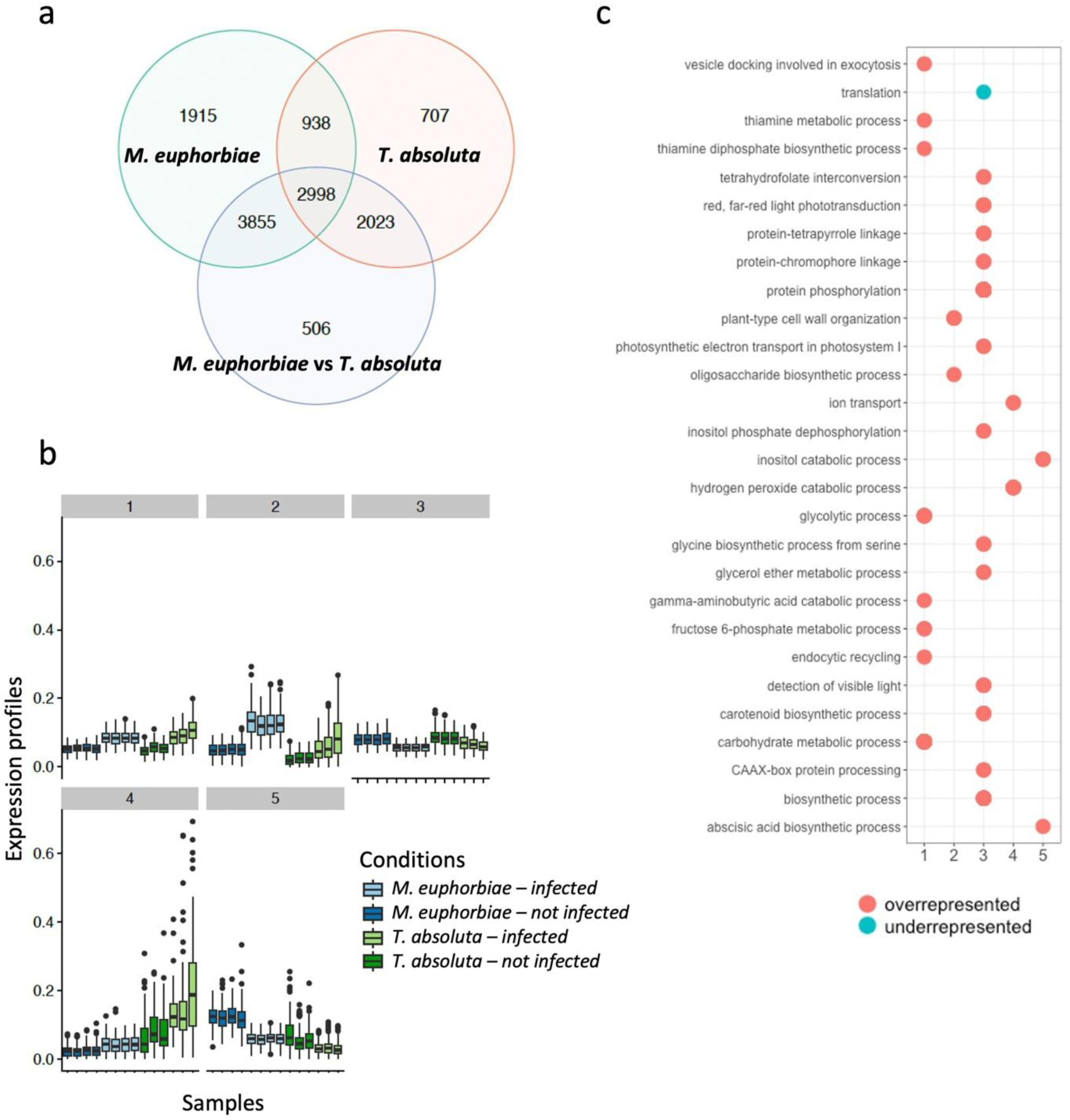
- *S. lycopersicum* leaf response to *M. euphorbiae* and *T. absoluta* highlighted 938 genes displaying a similar response to both parasites. (a) Venn diagram of differentially expressed genes in *S. lycopersicum* leaves in response to *M. euphorbiae (in green)* and *T. absoluta (in red) and the genes significantly differ in their response between the two parasites (M. euphorbiae vs T. absoluta in blue*) based on generalized linear model and likelihood ratio test-based p-values adjusted lower than 0.05 (Benjamini-Hochberg) based on the EdgeR package (Robinson et al., 2010) via DiCoExpress (Lambert et al., 2020). (b) Co-expression clusters with the boxplot expression profiles per sample per conditions in leaves. (c) GO enrichment of biological processes per co-expression cluster gene candidates commonly dysregulated in presence of leaf pathogens.

We first focused on *S. lycopersicum* root response in presence of telluric parasites (*M. incognita* and *P. parasitica*). Multifactorial analysis of the RNA-seq data from the root parasite experiments resulted in 7,734 candidate genes significantly dysregulated in presence of *M. incognita* and *P. parasitica* among which 2,119 candidate genes showed a common response to root pathogens, (Figure 4a). Co-expression analysis focusing on these 2,119 candidates resulted in five co-expression clusters (Figure 4b and Figure S4b). Clusters 1, 2 and 3 contained genes generally up-regulated in response to both root pathogens (*M. incognita* and *P. parasitica*), whereas clusters 4 and 5 were characterized by a general pattern of downregulation in response to root pathogens. We identified over-represented cluster-specific biological processes using a GO enrichment analysis for each cluster (Figure 4c). Cluster 1 was characterized by an overrepresentation of processes linked with the regulation of SAR and ethylene-activated signaling pathway, including the candidate hub NIM1-INTERACTING 2-like (*Solyc04g007140*) (Figure 1f). In contrast, clusters 2 and 3 showed an enrichment in processes associated with programmed cell death (PCD) and regulation of jasmonic acid mediated signaling pathway (including the NIM1-like protein 1 encoded by *Solyc02g069310*) (Figure 3a). Both clusters were also enriched for GO term “regulation of SA mediated signaling pathway”. Processes linked to trehalose biosynthetic process and response to hormone were overrepresented in clusters 4 and 5, respectively (Figure 4c). Notably, clusters 3 and 4 shared an enrichment in processes linked to protein phosphorylation despite displaying opposite co-expression patterns. This suggests that genes belonging to the same pathway may have opposite roles (positive /negative regulators), and that parasites may need to up-regulate some components while downregulating others to achieve parasitic success (Figure 4c).

We then performed a similar analysis with leaf pathogens (*M. euphorbiae* and *T. absoluta*). Multifactorial analysis identified 938 genes showing a common response to leaf pathogens (Figure 5a). Co-expression analysis of these candidates also resulted in five co-expression clusters with clusters 1,2 and 4 containing genes generally up-regulated in response to both leaf parasites (*M. euphorbiae* and *T. absoluta*), whereas clusters 3 and 5 were characterized by a general pattern of downregulation in response to leaf parasites (Figure 5b and Figure S4c). As observed with root parasites, GO enrichments were cluster specific (Figure 5c). Cluster 1 showed an overrepresentation of processes related to vesicle trafficking and carbohydrate metabolic process, with candidates associated with cell wall remodeling such as the beta-mannosidase (*Solyc03g119080*) (Pirona et al., 2023). Additional cell wall-associated genes, such as *Solyc09g098510* (extensin-2-like), were present in cluster 2, which was enriched in plant-type cell wall organization processes. Cluster 3 displayed enrichments in processes associated with protein phosphorylation and photosynthesis. Among the genes in cluster 3, the MAP kinase kinase kinase encoding gene *Solyc02g065110* has been associated with the induction of PCD (Roberts et al., 2019). Cluster 4 was enriched in ion transport processes with cyclic nucleotide-gated ion channel candidates, consistent with the changes in ion fluxes associated with plant immune response to pathogens (Moeder et al., 2011) and in hydrogen peroxide catabolic process. Finally, cluster 5 was enriched in inositol catabolism with downregulated inositol oxygenases, a gene family involved in the cell wall biosynthesis and ascorbic acid biosynthesis pathway (Munir et al., 2020). Together, these results indicate that distinct leaf parasites trigger a common transcriptional response involving coordinated modulation of cell wall dynamics, vesicle trafficking, ion homeostasis, and photosynthesis. Although several candidate genes responded to both root and leaf pathogens, the co-expression analyses revealed a panel of biological processes significantly over-represented in a cluster and tissue specific manner. These results unveiled clear distinct modules of underlying regulations associated with pathogen responses.

### Putative *S. lycopersicum* hubs targeted by parasites are homologs of *A. thaliana* hubs

Finally, we evaluated the relevance of the 1,606 candidate hubs commonly up or down regulated in *S. lycopersicum* in response to *M. incognita*, *P. parasitica* and *M. euphorbiae* (Figure 1c) within the experimentally tested *A. thaliana* interactome network available on BioGRID (Stark et al., 2006) in a model-to-crop translational work using OrthoFinder-identified orthologs as proxies. We characterized over 900 homologs of the putative hubs identified in this study within the *A. thaliana* interactome network, most of which taking part in complex, highly interconnected networks (Figure 6a, Figure S5 and Table S1). To reconstruct the networks affected by the three parasites, we then considered only the interactions between the homologs of our candidate hubs. This analysis revealed the strategic positions of several effector-interacting homologs, such as the PP2CA (Solyc03g096670 ortholog, Degree = 13 and Betweenness Centrality = 0.092) and the plasma membrane intrinsic protein PIP2A (associated with paralogs Solyc06g011350, Solyc09g007760 and Solyc09g007770, Degree = 24 and Betweenness Centrality = 0.262), among numerous other candidates (Figure 6b and Table S2). Further focus on *A. thaliana* homologs of the *S. lycopersicum* candidate genes displaying a log2FC > 2 in response to *M. incognita*, *P. parasitica* and *M. euphorbiae* pinpointed the pivotal position of NATA1 (AT2G39030, Solyc08g006765 homolog) (Figure 6c and Table S3). NATA1 interacts with two transcription factors, HB-12 (Solyc01g096320 homolog) and TCP2 (Solyc08g048370 homolog), thereby bridging two important regulatory network subsets (NATA1 Degree = 2 and Betweenness Centrality = 0.158). AtHB12 acts as positive transcriptional regulators of *PP2C* genes, and thus as a negative regulator of ABA signaling (Valdés et al., 2012). On the other branch of this network, we identified three TCP transcription factors, TCP2, TCP21 and TCP24, which centralize interactions involving numerous differentially expressed genes identified in our study. TCP2 and TCP21 are known to be directly targeted by pathogen’s effectors (González-Fuente et al., 2020; Drcelic et al., 2024; Fu et al., 2025). Consistently, in our study, Solyc08g048370 and Solyc01g008230, the tomato homologs of TCP2 and TCP21, were similarly dysregulated in response to the three parasites, making them strong candidate hubs (Figure S6).

**Figure 6.**
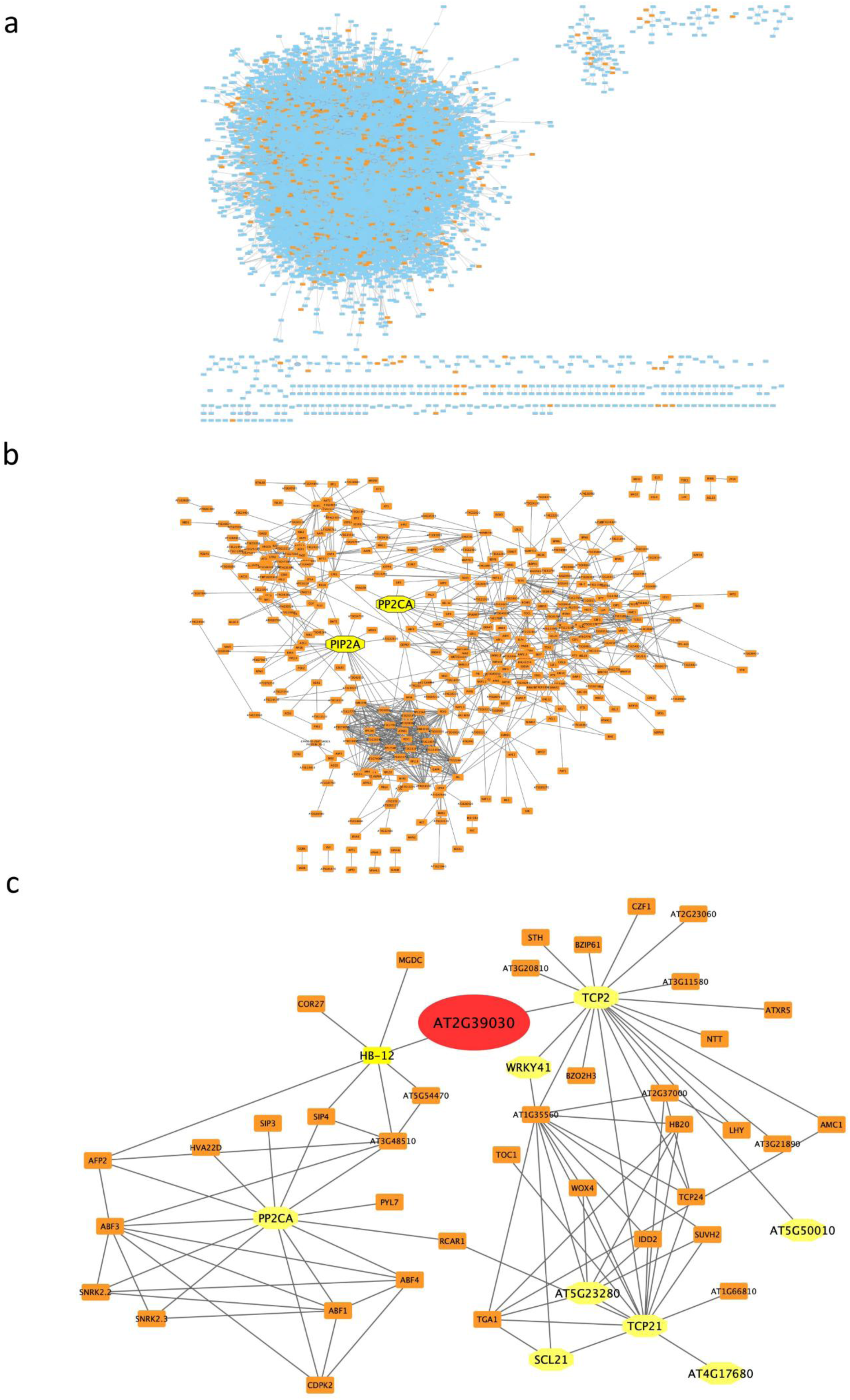
- S. lycopersicum orthologs in the context of experimentally tested A. thaliana regulatory network. (a) BioGRID (Stark et al., 2006) based A. thaliana full interactome network on which the orthologs/homologs of the 1,606 candidate hubs commonly up or down-regulated in S. lycopersicum in response to M. incognita, P. parasitica and M. euphorbiae were highlighted in orange. (b) A. thaliana interactome subset showing only the interactions between the orthologs/homologs of the 1,606 candidate hubs commonly up or down-regulated in S. lycopersicum in response to M. incognita, P. parasitica and M. euphorbiae (Degree > 0). (c) A. thaliana interactome subset highlighting NATA1 (AT2G39030 in red) and its interactants among the orthologs/homologs of candidate hubs commonly up or down-regulated in S. lycopersicum in response to M. incognita, P. parasitica and M. euphorbiae. In yellow: effector interacting proteins from EffectorK (González-Fuente et al., 2020).

## Discussion

Compatible plant-pathogen interactions rely on the pathogens ability to manipulate host gene expression in order to (1) facilitate pathogen entry through interference with the integrity of physical barriers such as modulations of the cell wall structure or stomatal opening, (2) suppress host immune responses by altering gene expression regulation and hormonal signaling pathways and (3) promote pathogen access to host nutrients via the regulation of metabolite synthesis and nutrient transport (van Schie & Takken, 2014; D. Wang et al., 2025). Such genes can be evolutionary conserved across species (Koseoglou et al., 2022) and may be targeted, directly or indirectly, by effectors from trans-kingdom pathogens. Understanding these hub targets in depth could therefore contribute to the development of broad-spectrum resistance through their targeted impairment.

In this context, we performed a study dissecting tomato response in presence of three distinct parasites (nematodes, oomycetes and aphids). Although pathogens generally downregulate defense pathways while activating processes that favor their own growth, host responses reflect complex regulatory networks with redundancy and feedback loops, complicating transcriptomic interpretation. Illustrating this conundrum, we found that *Solyc08g080670* was among the most strongly upregulated genes in response to all three parasites tested. This gene is the tomato homolog of *OSM34* in *A. thaliana,* a stress-responsive gene, functioning in the ABA signaling pathway, and potentially playing a role in plant protection against fungi infections (Mukherjee et al., 2010; Hao et al., 2015; Park & Kim, 2021). A recent study demonstrated that the TuMV hijacked *AtOSM34* to promote viral infection. Indeed, *AtOSM34* upregulation reduces plasmodesmata callose deposition to promote viral intercellular movement and targeted viral replication complex to promote viral replication. Additionally, it suppressed the ROS-mediated antiviral response (He et al., 2025). Whether multiple pathogens trigger the upregulation of *Solyc08g080670* to promote infection remains to be determined, but this possibility highlights its potential importance as a key factor to target. Another set of key candidate genes identified in this study are *SlDMR6-1* and *SlDMR6-2,* the closest homologs of the salicylic Acid 5-Hydroxylase DMR6 and DLO, which are well-known susceptibility factors to oomycetes and bacteria. In *Arabidopsis*, these enzymes function redundantly, in different tissues, to suppress plant immunity. They catalyze the formation of 2,5-Dihydroxybenzoic acid2 (5-DHBA) by hydroxylating SA, thereby inactivating SA (Zeilmaker et al., 2015; van Butselaar et al., 2024). Here, we showed that both genes are strongly overexpressed in tomato in response to all three pathogens tested. Interestingly, *SlDMR6-1* is only slightly induced in response to aphids, consistent with the variable spatial expression mentioned in the literature and with the possibility that *SlDMR6-1* and SlDMR6-2 respond differently depending on the pathogen. For instance, Thomazella et al. (2021) reported an upregulation of *SlDMR6-1* whereas SlDMR6-2 expression was not detected upon infection by *Xanthomonas gardneri*(Thomazella et al., 2021). Although such genes represent potential targets for gene inactivation to enhance plant resistance, their role in maintaining SA homeostasis highlights the trade-off between immunity and growth.

Another important multi-pathogen target identified in our study is the *NIMIN-2* homolog (Solyc04g007140). NIMIN proteins interact with NPR1, the key positive regulator of SAR acting in the SA-mediating pathway (Zavaliev et al., 2020). Interestingly, NIMIN proteins act as negative regulators of NPR1, therefore, the upregulation of *Solyc04g007140* is likely to favor pathogen infection by compromising SAR, making it a prominent target (Weigel et al., 2005). Pathogen infections also affected host epigenetic factors. Indeed, *Solyc06g066420* belongs to the *OXS 3* family, which, like OSM34, functions in ABA signaling and acts as a chromatin-associated factor involved in tolerance to heavy metals and oxidative stress (Blanvillain et al., 2009). *AtOXS3* has been associated with the regulation of gene expression via modulation of histone γ-H2A.X occupancy (Xiao et al., 2021). *Solyc06g066420* was consistently upregulated across all interactions and is therefore likely to modulate a wide range of gene expression programs. Conversely, *Solyc05g052950*, *AtTCF1* ortholog, was strongly downregulated. AtTCF1 has been studied in the context of abiotic stress, particularly cold acclimatation (Ji et al., 2015). TCF1 indirectly affects lignin accumulation though its association with chromatin at specific loci. Interestingly, loss of TCF1 modified H3K4me2 and H3K27me3 levels, resulting in reduced lignin content (Ji et al., 2015). Because lignin-based barrier restricts pathogen spread and contributes to plant resistance (Lee et al., 2019), *Solyc05g052950* is therefore an excellent epigenetic factor to target, acting upstream lignin accumulation. Together, these findings underscore the importance of chromatin-level regulation during parasitism. Furthermore, several candidate hubs identified here are also associated with abiotic stress responses (Pirona et al., 2023) reflecting conserved physiological responses—such as ABA signaling, lignin deposition, and stomatal regulation—between biotic and abiotic stress adaptation. This overlap highlights the potential cost of manipulating such hubs for resistance breeding.

Besides, subtilases are involved in a broad range of biological functions including plant defense responses or in plant-pathogen interactions (Weng et al., 2014; Serrano et al., 2016; Stegmann et al., 2017; J. Figueiredo et al., 2018; Zhang et al., 2024). Within this large gene family (around one hundred in *S. lycopersicum*), we identified two genes (from the SBT1.9 clade) that were strongly upregulated in response to all three pathogens, while 65 tomato subtilase encoding genes were differentially expressed in response to at least one of them. This result suggests that subtilases are primarily involved in pathogen-specific plant responses. In *S. lycopersicum*, research has mainly focused on the subtilase cluster encoding the P69 proteases, which has been linked to pathogenesis (Jordá et al., 1999; Zhang et al., 2024). For instance, *Pseudomonas syringae* and *Ralstonia solanacearum* infections have been associated with the induction of *P69B* among others (Jordá et al., 1999; Zhang et al., 2024). In our study, *P69B* (*Solyc08g079870*) was also up-regulated in response to *M. incognita*, *P. parasitica* and *M. euphorbiae*. In contrast, little is known about the role of the subtilases of the cluster 1, highlighting an important avenue for future studies to determine their role in pathogenesis. SBT1.9-related sequences (At5g67090-related) have undergone a large expansion in tomato, which raises the question of their putative function. Besides a hypothetical neofunctionalization of these candidates, the large number of SBT1.9-like members in tomato may be due to the diversity of their substrates in tomato. Indeed, proteases such as caspase in animals or phytaspase in plants, recognize several amino acid residues of substrates, and this recognition can be highly specific, particularly in critical pathways such as programmed cell death. Thus, SBT1.9-related sequences would ensure a highly specific recognition for several pivotal substrates to process that still need to be uncovered.

SA-activated responses are effective against biotrophic pathogens, which retrieve nutrients from living plant cells. In contrast, JA and ET elicit responses that combat necrotrophic microbes, which kill plant cells and feed on the resulting tissues. While activation of the JA pathway suppresses SA biosynthesis and vice versa (Zheng et al., 2012), SA signaling also interferes with JA-ET pathways.

For example, SA affects JA-induced transcription by inducing the degradation of transcription factors that activate JA signaling, as shown for the ERF transcription factor ORA59. Additionally, SA can induce negative regulators, among them several WRKY transcription factors, that directly or indirectly inhibit JA-responsive gene expression (Zheng et al., 2012). More than one hundred transcription factors were commonly dysregulated in response to the three pathogens tested. We observed a divergence in the TF family enrichments between the up and down regulated candidates. Similar results were also observed in transcriptomic analysis of *S. lycopersicum* in response to osmotic stress (Pirona et al., 2023). Interestingly, our results and those of (Pirona et al., 2023) showed that, in response to both biotic and abiotic stresses, ERF and bHLH families of transcription factors were among the top enrichment in up and down regulated candidates, respectively, with several common candidate genes responding to both types of stresses.

GO enrichment analyses revealed common overrepresentation of metabolic processes across nematode, aphid, and oomycete infections, supporting the view that parasites enhance host metabolic activity. Beyond a set of commonly dysregulated candidate hubs, roots and leaves diverged markedly in the significantly overrepresented processes identified across the different co-expression clusters. These findings, along with previous studies reporting tissue-specific responses between leaves and roots (Hermanns et al., 2003; Attard et al., 2010), indicate that combining organ-specific with common hubs may be a more effective strategy for engineering broad-spectrum resistance. Such an approach could be implemented through rootstock–scion combinations, in which each organ is optimized according to its specific functional requirements. Candidate hubs could therefore be selected based on their tissue specificity, with root-targeted hubs engineered in the rootstock and leaf- or shoot-specific hubs in the scion. By combining complementary regulatory networks within a single plant, this strategy could generate finely tuned genotypes that optimize plant fitness, crop yield, and enhance durable resistance against a broad spectrum of pests.

Finally, we identified *Solyc08g006765*, the tomato homolog of *AtNATA1*, as a central multi-pathogen hub. Insect feeding and *P. syringae* inoculation both upregulate NATA1, leading to the production of *N*^δ^-acetylornithine via the jasmonate pathway (Adio et al., 2011). While this metabolite affects the reproduction of the aphid, *Myzus persicae*, *P. syringae* growth was impaired in *nata1* mutant. In addition to jasmonic acid, the coronatine effector produced by DC3000 *P. syringae* strain also induces *NATA1* expression (Lou et al., 2016). Via the acetylation of putrescine, a precursor required for the defense-related hydrogen peroxide, NATA1 acts as a susceptibility factor competing for this common substrate impeding plant defense response (Lou et al., 2016). Furthermore, we noted that several NATA1 interactors, including TCP transcription factors, are also direct targets of known effectors (González-Fuente et al., 2020; Drcelic et al., 2024; Fu et al., 2025). We identified tomato homologs of AtTCP2 and AtTCP21 as consistently dysregulated in presence of all three pathogens. Although *SlTCP2* and *SlTCP21* belong to different classes and are probably involved in two different pathways, their expression is clearly affected by multiple pathogens. Additionally, TCP21, also named CHE for CCA1 Hiking Expedition, was recently described to be the missing link between local infection and the SAR making it a compelling hub (Cao et al., 2024). We propose that these transcription factors and especially *TCP21* (*CHE*), should be considered as a key regulatory hub in future studies of tomato pathogen interactions.

Far beyond the trivial production of a repertoire of commonly dysregulated genes, we implemented an in-depth co-expression analysis that pinpoint compelling candidate hubs covering a large range of functions and pathways. By pairing our results obtained in tomato with *Arabidopsis* interactome network data, we found crucial factors associated with parasitic success that should be of interest for plant breeding programs.

## Acknowledgments

The authors are grateful to Marie-Laure Martin and Etienne Delannoy (IPS2, Orsay, France) for their guidance with the co-expression analysis and Romain Larbat (IRHS, Angers, France) for sharing the RNA-seq datasets of tomato response to *T. absoluta* (Roumani et al., 2022). The authors thank the bioinformatics and genomics platform BIG (ISA, Sophia Antipolis, France, plantBIOs, https://doi.org/10.15454/qyey-ar89) for computing and storage resources and the Genotoul bioinformatics platform (Bioinfo Genotoul, Toulouse, France https://doi.org/10.15454/1.5572369328961167E12) for computing resources.

## Data, scripts, code, and supplementary information availability

The datasets generated for this study are available in the NCBI BioProject repository under the accession number PRJNA1375716. The datasets of tomato response to *T. absoluta* infection at 24 hours post infection (Roumani et al., 2022) were retrieved through the accession number GSE200795. Supplementary information and code are available at https://doi.org/10.57745/GL5MLM and https://forge.inrae.fr/GNet/dicoexpress/-/tree/985351e4e49c854f1bcb894de585c6939374f8bc/ and https://forge.inrae.fr/plantbios_big/nextflow/mapping_count_featurecount_rsem/-/tree/f9f4b7dd59fb5cc295c446ab9fd067a41956071c/, respectively.

