## Supplemental Figures for "Transcriptional responses of *Solanum lycopersicum* to three distinct parasites reveal host hubs and networks underlying parasitic successes"

**Affiliations**

***Supplementals***

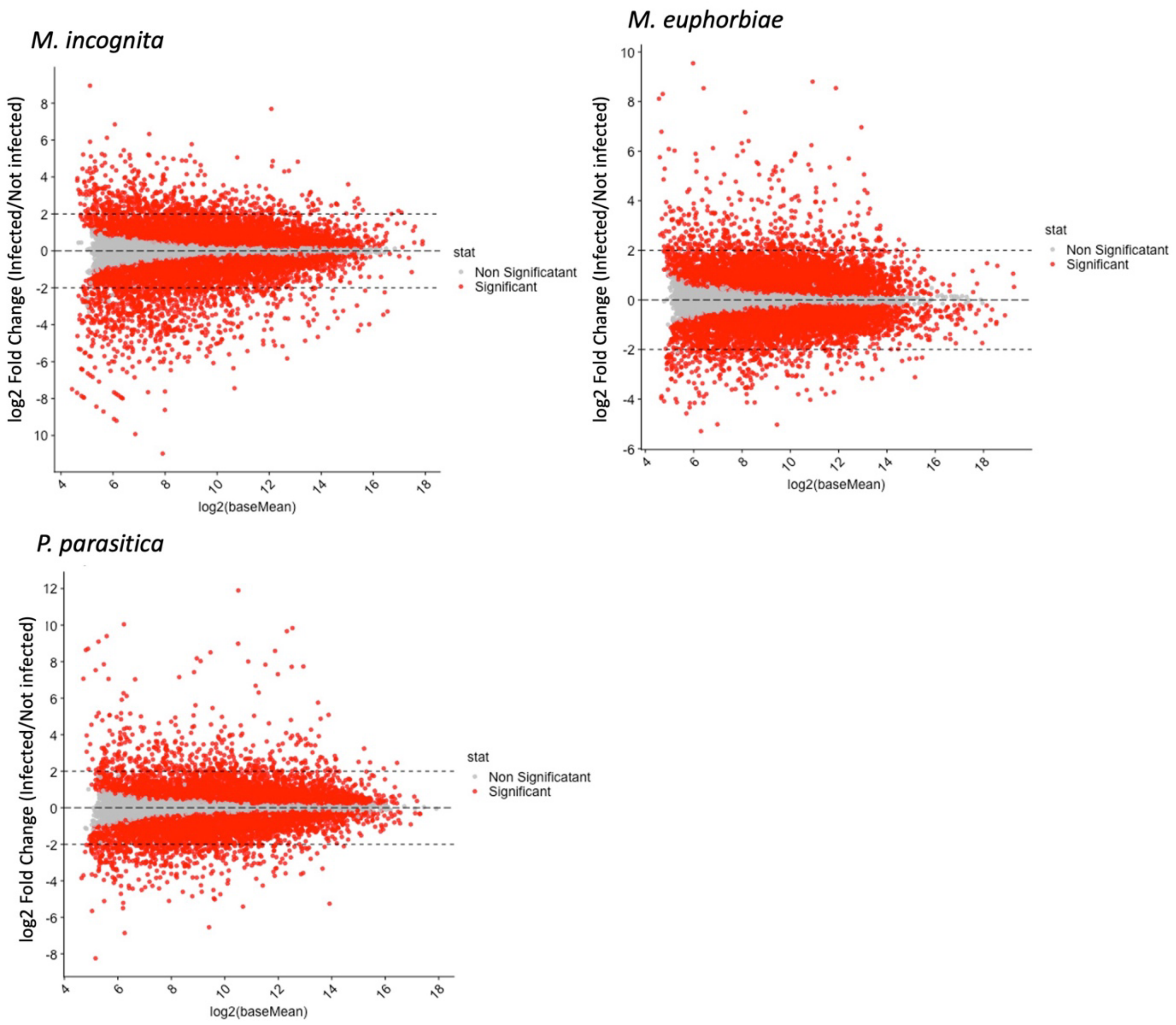


*Figure S1: Minus-Average (MA) plots showing the change in gene expression in S. lycopersicum in response to parasite attacks (M. incognita, M. euphobiae and P. parasitica) vs non-infected samples (DESeq2 (Love et al., 2014), adjusted p-value lower than 0.05 - FDR/Benjamini-Hochberg, log2 Fold Change (Infected/Not infected)).*

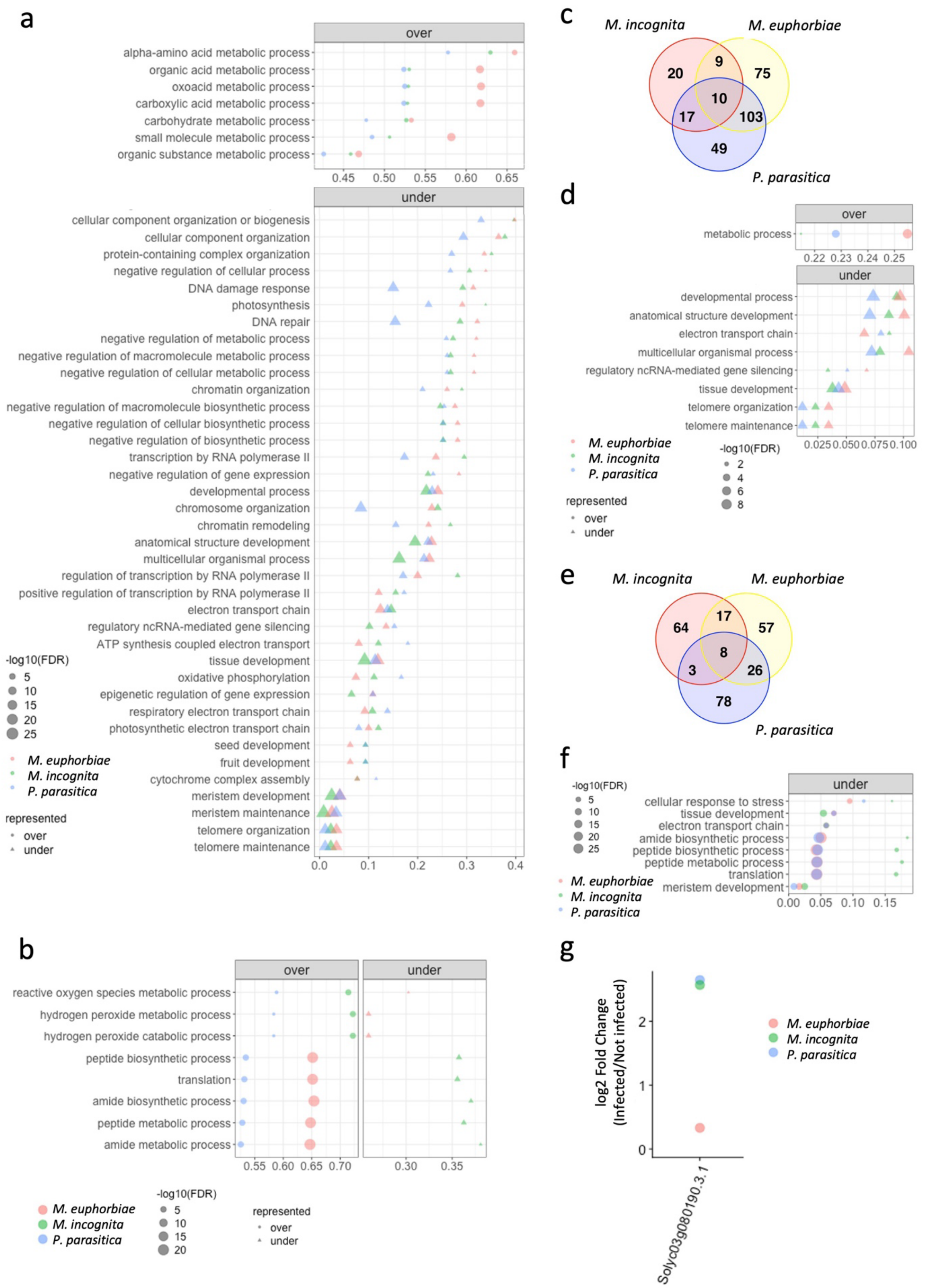


*Figure S2: Biological processes GO term enrichment analysis of S. lycopersicum response to M. euphorbiae, M. incognita and P. parasitica. a. GO terms commonly over or under represented in all three interactions (M.*

*euphorbiae, M. incognita and P. parasitica); b. GO terms in common between the three interactions which were either over or under represented depending on the interaction. c. Venn Diagrams of GO overlaps between the three interactions considering only the differentially expressed genes up-regulated in response to at least one parasite. d. GO terms commonly over or under represented in all three interactions (M. euphorbiae, M. incognita and P. parasitica) considering only the differentially expressed genes up-regulated in response to at least one parasite. e. Venn Diagrams of GO overlaps between the three interactions considering only the differentially expressed genes down-regulated in response to at least one parasite. f. GO terms commonly over or under represented in all three interactions (M. euphorbiae, M. incognita and P. parasitica) considering only differentially expressed genes down-regulated in response to at least one parasite. g. SlDMR6-1 (Solyc03g080190) expression profiles in response to M. euphorbiae, M. incognita and P. parasitica. (DESeq2 (Love et al., 2014), adjusted p-value lower than 0.05 - FDR/Benjamini-Hochberg, log2 Fold Change (Infected/Not infected)).*

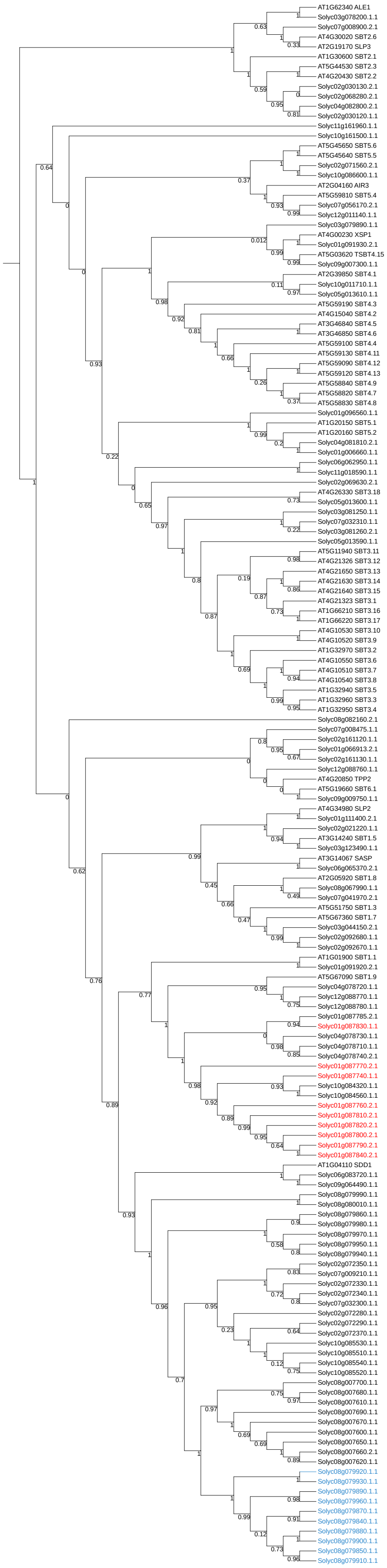
AT1G62340 ALE1
Solyc03g078200.1.1
Solyc07g008900.2.1
AT4G30020 SBT2.6
AT2G19170 SLP3
AT1G30600 SBT2.1
AT5G44530 SBT2.3
AT4G20430 SBT2.2
Solyc02g030130.2.1
Solyc02g068280.2.1
Solyc04g082800.2.1
Solyc02g030120.1.1
Solyc11g161960.1.1
Solyc10g161500.1.1
AT5G45650 SBT5.6
AT5G45640 SBT5.5
Solyc02g071560.2.1
Solyc10g086600.1.1
AT2G04160 AIR3
AT5G59810 SBT5.4
Solyc07g056170.2.1
Solyc12g011140.1.1
Solyc03g079890.1.1
AT4G00230 XSP1
Solyc01g091930.2.1
AT5G03620 TSBT4.15
Solyc09g007300.1.1
AT2G39850 SBT4.1
Solyc10g011710.1.1
Solyc05g013610.1.1
AT5G59190 SBT4.3
AT4G15040 SBT4.2
AT3G46840 SBT4.5
AT3G46850 SBT4.6
AT5G59100 SBT4.4
AT5G59130 SBT4.11
AT5G59090 SBT4.12
AT5G59120 SBT4.13
AT5G58840 SBT4.9
AT5G58820 SBT4.7
AT5G58830 SBT4.8
Solyc01g096560.1.1
AT1G20150 SBT5.1
AT1G20160 SBT5.2
Solyc04g081810.2.1
Solyc01g006660.1.1
Solyc06g062950.1.1
Solyc11g018590.1.1
Solyc02g069630.2.1
AT4G26330 SBT3.18
Solyc05g013600.1.1
Solyc03g081250.1.1
Solyc07g032310.1.1
Solyc03g081260.2.1
Solyc05g013590.1.1
AT5G11940 SBT3.11
AT4G21326 SBT3.12
AT4G21650 SBT3.13
AT4G21630 SBT3.14
AT4G21640 SBT3.15
AT4G21323 SBT3.1
AT1G66210 SBT3.16
AT1G66220 SBT3.17
AT4G10530 SBT3.10
AT4G10520 SBT3.9
AT1G32970 SBT3.2
AT4G10550 SBT3.6
AT4G10510 SBT3.7
AT4G10540 SBT3.8
AT1G32940 SBT3.5
AT1G32960 SBT3.3
AT1G32950 SBT3.4
Solyc08g082160.2.1
Solyc07g008475.1.1
Solyc02g161120.1.1
Solyc01g066913.2.1
Solyc02g161130.1.1
Solyc12g088760.1.1
AT4G20850 TPP2
AT5G19660 SBT6.1
Solyc09g009750.1.1
AT4G34980 SLP2
Solyc01g111400.2.1
Solyc02g021220.1.1
AT3G14240 SBT1.5
Solyc03g123490.1.1
AT3G14067 SASP
Solyc06g065370.2.1
AT2G05920 SBT1.8
Solyc08g067990.1.1
Solyc07g041970.2.1
AT5G51750 SBT1.3
AT5G67360 SBT1.7
Solyc03g044150.2.1
Solyc02g092680.1.1
Solyc02g092670.1.1
AT1G01900 SBT1.1
Solyc01g091920.2.1
AT5G67090 SBT1.9
Solyc04g078720.1.1
Solyc12g088770.1.1
Solyc12g088780.1.1
Solyc01g087785.2.1
Solyc01g087830.1.1
Solyc04g078730.1.1
Solyc04g078710.1.1
Solyc04g078740.2.1
Solyc01g087770.2.1
Solyc01g087740.1.1
Solyc10g084320.1.1
Solyc10g084560.1.1
Solyc01g087760.2.1
Solyc01g087810.2.1
Solyc01g087820.2.1
Solyc01g087800.2.1
Solyc01g087790.2.1
Solyc01g087840.2.1
AT1G04110 SDD1
Solyc06g083720.1.1
Solyc09g064490.1.1
Solyc08g079990.1.1
Solyc08g080010.1.1
Solyc08g079860.1.1
Solyc08g079980.1.1
Solyc08g079970.1.1
Solyc08g079950.1.1
Solyc08g079940.1.1
Solyc02g072350.1.1
Solyc07g009210.1.1
Solyc02g072330.1.1
Solyc02g072340.1.1
Solyc07g032300.1.1
Solyc02g072280.1.1
Solyc02g072290.1.1
Solyc02g072370.1.1
Solyc10g085530.1.1
Solyc10g085510.1.1
Solyc10g085540.1.1
Solyc10g085520.1.1
Solyc08g007700.1.1
Solyc08g007680.1.1
Solyc08g007610.1.1
Solyc08g007690.1.1
Solyc08g007670.1.1
Solyc08g007600.1.1
Solyc08g007650.1.1
Solyc08g007660.2.1
Solyc08g007620.1.1
Solyc08g079920.1.1
Solyc08g079930.1.1
Solyc08g079890.1.1
Solyc08g079960.1.1
Solyc08g079870.1.1
Solyc08g079840.1.1
Solyc08g079880.1.1
Solyc08g079900.1.1
Solyc08g079850.1.1
Solyc08g079910.1.1

*Figure S3: Extended phylogenetic tree of subtilase family from S. lycopersicum and A. thaliana. The Maximum Likelihood phylogenetic tree (WAG model, aLRT test) contains 55 subtilases from A. thaliana and 103 subtilases from S. lycopersicum. Cluster 1 from S. lycopersicum in red and Cluster 8 from S. lycopersicum in blue.*

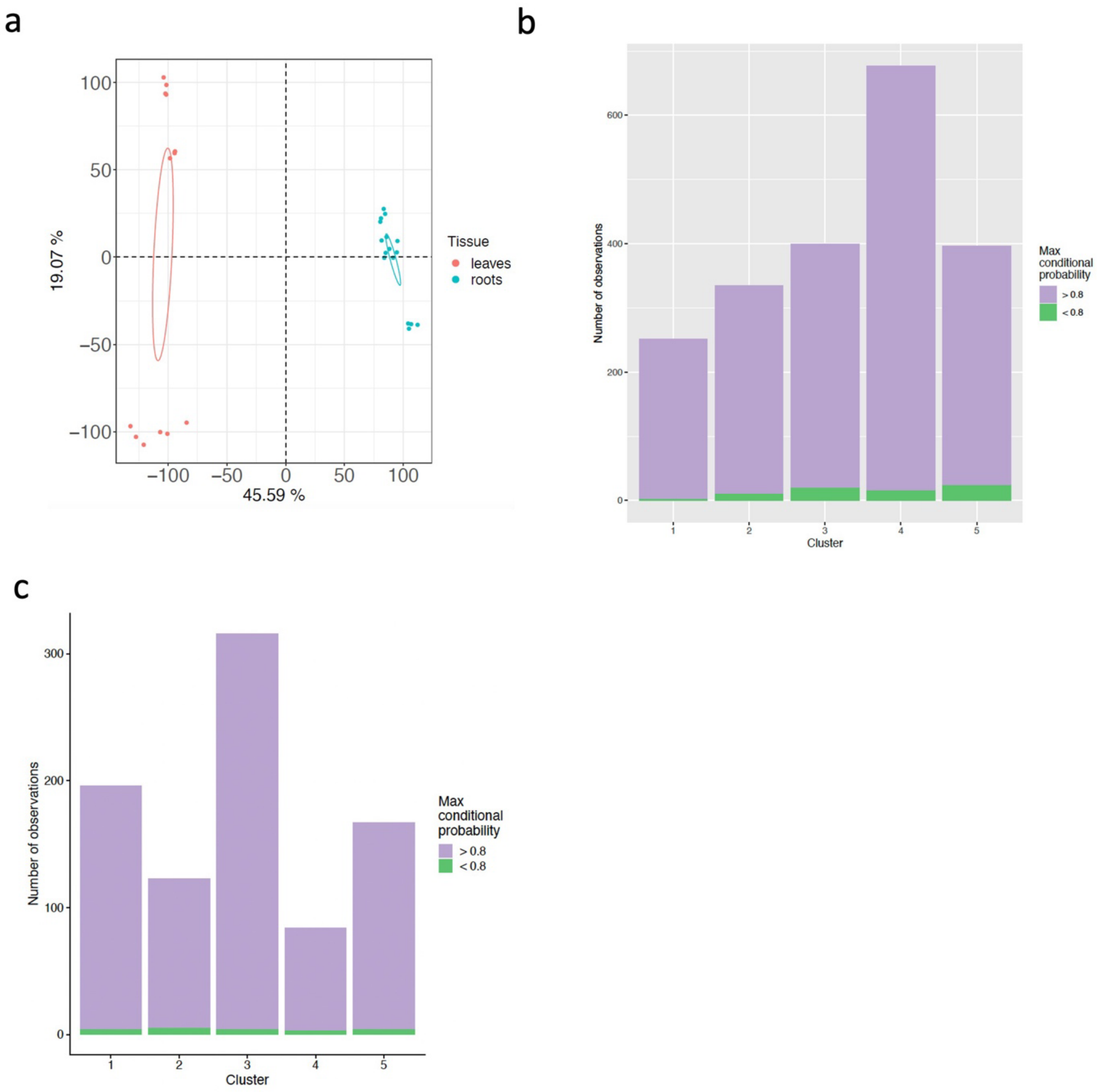


*Figure S4: Multifactorial RNA-seq analysis of S. lycopersicum response to M. incognita, P. parasitica, M. euphorbiae and T. absoluta using DiCoExpress (Lambert et al., 2020). a. Principal component analysis (PCA) of S. lycopersicum responses to root (M. incognita and P. parasitica) and leaf (M. euphorbiae and T. absoluta) parasites based on the RNA-seq samples showing a clear separation between leaf and root samples along the first principal component. b. Max conditional probabilities associated with each co-expression cluster (1-5) identified in Figure 4b based on S. lycopersicum root response to telluric parasites (M. incognita and P. parasitica) using DiCoExpress (Lambert et al., 2020). c. Max conditional probabilities associated with each co-expression cluster (1-5) identified in Figure 5b based on S. lycopersicum leaf response to M. euphorbiae and T. absoluta using DiCoExpress (Lambert et al., 2020).*

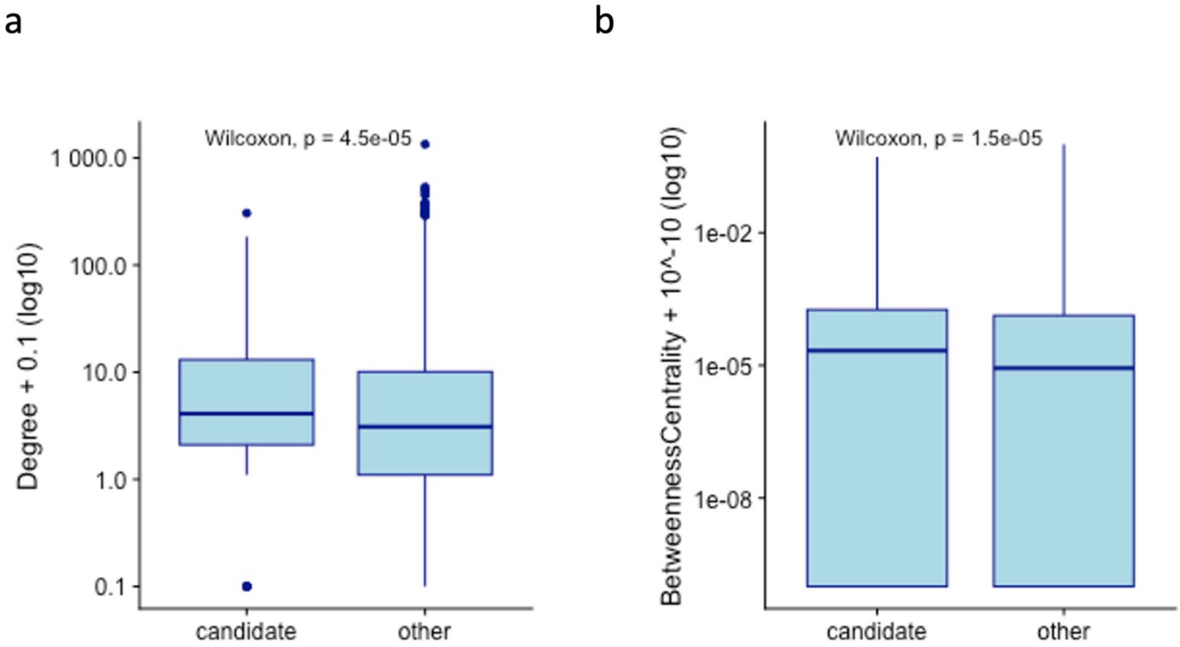


*Figure S5: Degree and Betweenness Centrality characteristics of the orthologs/homologs of candidate hubs identified in S. lycopersicum within the A. thaliana interactome network. a. Degree distributions in candidates vs non candidates within the A. thaliana interactome network related to Figure 5a. b. Betweenness Centrality distributions within the A. thaliana interactome network related to Figure 5a.*

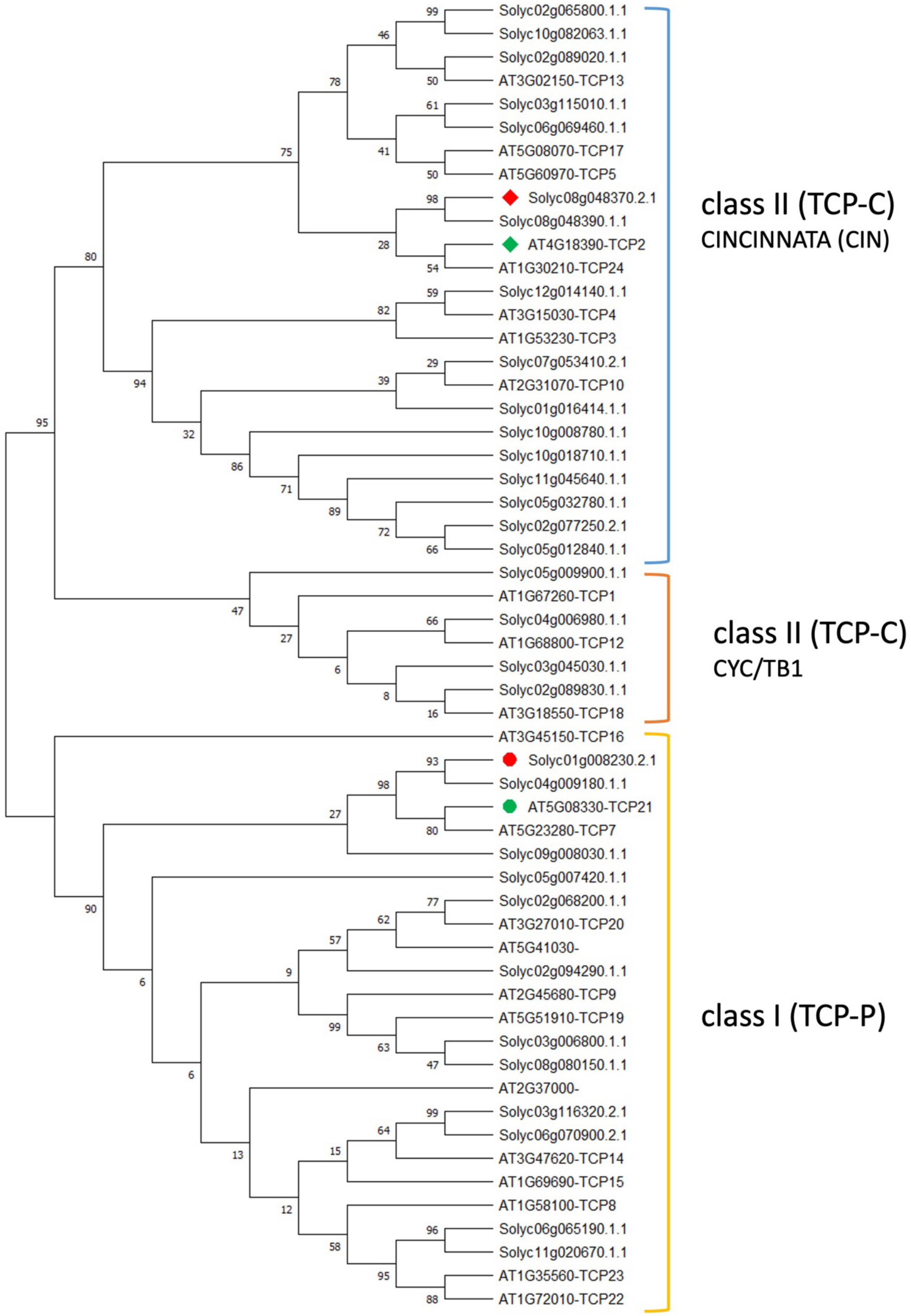


*Figure S6: Phylogenetic tree showing the 24 and 32 proteins annotated as TCP transcription factors from A. thaliana and S. lycopersicum. Accession names are related to the protein database used, Araport11 and ITAG4.1 (AT for A. thaliana and Solyc for S. lycopersicum). The evolutionary history was inferred using the Neighbor-Joining method. The bootstrap consensus tree was inferred from 1000 replicates. The evolutionary distances were computed using the JTT matrix-based method. TCP sub-classes are indicated.*

Love MI, Huber W, Anders S (2014) Moderated estimation of fold change and dispersion for RNA-seq data with DESeq2. *Genome Biol,* **15**, 550. https://doi.org/10.1186/s13059-014-0550-8
